# The Kishony Mega-Plate Experiment, *a Markov Process*

**DOI:** 10.1101/2021.12.23.474071

**Authors:** Alan Kleinman

## Abstract

A correct understanding of the DNA evolution of drug resistance is critical in developing strategies for suppressing and preventing this process. The Kishony Mega-Plate Experiment demonstrates this important phenomenon that occurs in the practice of medicine, that of the evolution of drug-resistance. The evolutionary process which the bacteria in this experiment are doing is called a Markov Process or Markov Chain. Understanding this process enables clinicians and researchers to predict the evolution of drug-resistance and develop strategies to prevent this process. This paper will show how to apply the Markov Chain model of DNA evolution to the Kishony Mega-Plate Experiment and why the experiment behaves the way it does by contrasting the Jukes-Cantor model of DNA evolution (a stationary model) with a modification of the Jukes-Cantor model that makes it a non-stationary, non-equilibrium Markov Chain. The numerical behaviors of the stationary and non-stationary models are compared. What this analysis shows is that DNA evolution is a non-stationary, non-equilibrium process and that by using the correct non-stationary, non-equilibrium model that one can simulate and predict the behavior of real evolutionary examples and that these analytical tools can give the clinician guidance on how to use antimicrobial selection pressures for treating infectious diseases. This in turn can help reduce the numbers and costs of hospitalization for sepsis, pneumonia and other infectious diseases.

## Introduction

The Kishony Mega-Plate Experiment is described in the paper “Spatiotemporal microbial evolution on antibiotic landscapes” [1]. A description of how to make a Mega-Plate Experiment can be found in reference [2]. Many videos of the experiment can be found on the internet, a good example can be found here [3]. The next paragraph gives a simple explanation of the experiment.

The experiment consists of the following. A large petri dish is constructed. Growth media is placed on this petri dish, and in this growth media, different concentrations of an antimicrobial agent are placed in bands in the growth media. In the left and right-hand bands in the petri dish, no antimicrobial agent is used. In the next adjacent bands to these zero concentration bands, the lowest concentration of the drug is used. As one move to the center of the petri dish, increasing concentrations of the drug are placed in each band until in the middle of the dish that contains the highest concentration of the antimicrobial agent. Then a motile “wild-type” bacteria (founder) is introduced into the petri dish that has no resistance initially to the antimicrobial agent used and is only able to grow in the drug-free region. As these colonies in the drug-free regions grow, mutations occur in some members of these colonies. Occasionally, some member gets a beneficial mutation that enables it to grow in the next higher drug-concentration region. That new variant with the first resistance mutation now forms a new colony that can grow in the lowest drug concentration region. As that new colony grows, one of its members gets another beneficial mutation that allows it to grow in the next higher drug-concentration region. This process continues until finally there is a variant that can grow in the high drug-concentration region.

For this experiment to work in this size petri dish, the increase in concentration for adjacent bands must be limited so that it only requires a single beneficial mutation to occur on some member of the drug-sensitive variant population in the next lower drug-concentration band to grow in the next higher drug-concentration band. As the population increases in a particular band, descendants are getting mutations in their DNA as replications occur. This is a random walk process where the number of members taking their particular random walk increases as the population grows in number and as different variants occur. Some variants on their own particular random walk accumulate particular mutations that give improved fitness to the antibiotic selection pressure allowing these variants to grow in the higher drug-concentration regions. This process can be mathematically modeled as a Markov Process. A variety of different Markov Models of DNA evolution have been proposed. One of the earliest models is the Jukes-Cantor model presented in 1969 [4]. Many derivative models have been proposed such as the K80 [5] and K81 [6] (1980 and 1981 Kimura models respectively), the F81 Felsenstein model [7] presented in 1981, HKY85 model [8] (Hasegawa, Kishino and Yano 1985) model, the T92 model [9] (Tamura 1992), and the TN93 model [10] (Tamura and Nei 1993). The Jukes-Cantor and derivative models are commonly used in an attempt to compute the evolutionary distance between different sequences of homologous genetic code. Some recent examples of applications of the Jukes-Cantor and derivative models can be found in the following references, [11–15]. The main difference between the Jukes-Cantor model and the derivative models listed above is derivative models allow for different mutation rates for the different elements of the transition matrix, for example, to address the fact that base transversions can have a different mutation rate than for base transitions. However, the transition matrix is still constant over time.

Huelsenbeck and his co-authors wrote: “At present, a universal assumption of model-based methods of phylogenetic inference is that character change occurs according to a continuous-time Markov chain. At the heart of any continuous-time Markov chain is a matrix of rates, specifying the rate of change from one character state to another. For many phylogenetic analyses using DNA sequence data, it is assumed that there are four states (the nucleotides A, C, G, T/U) with a 4 × 4 matrix of rates among the 12 possible nucleotide substitutions. A few standard models of DNA substitution have been proposed. These include those first described by Jukes and Cantor (1969), Kimura (1980, 1981), Felsenstein (1981, 1984), Hasegawa, Yano, and Kishino (1984, 1985), Tamura and Nei (1993), and Tavare’ (1986).”[16] The intention here in this paper is to give a proposed modification of the heart of the Markov chain model that is demonstrated to correlate with the experimental model, the Kishony Mega-Plate experiment.

The Jukes-Cantor and derivative models listed above have a property in common. These models assume that the elements of the Markov transition matrix are constant with each replication. It is shown this implicit assumption gives a model constrained to constant population size, and that constant population size is 1. This assumption gives a model that fails to correctly model the evolution of antimicrobial resistance. A modification of the Jukes-Cantor model is presented that allows for modeling DNA evolution in different population sizes and that the population can vary at each evolutionary transitional step (replication). The mathematics for these two Markov models are described in the next section. The results are compared and the Markov model which takes into account population size is shown to compare favorably with the experimental results of the Kishony Mega-Plate experiment.

## Materials and Methods

### Statistical Analysis

The statistical analysis used in this study is based on the Markov process. A Markov process is a stochastic or random process where future outcomes can be predicted based solely on the present state of the system. A Markov process for discrete events is called a Markov chain. When formulated correctly, a Markov chain that models DNA evolution gives the theoretical distribution of frequencies of the different variants in a growing population based on the population size, the mutation rate, and the known initial state of the population. The formulation of such a model consists of the following steps.

The first step is to identify the possible states in which the system can be. When modeling DNA evolution, a given site in a genome can be in one of four possible states (one of the four possible DNA bases). When considering two sites simultaneously in a genome, those two sites can be in 16 possible states (4^2^ possible combinations of bases). When considering three sites, the number of possible states for the system is 64 possible states (4^3^ possible combinations). As one considers more sites in the genome, the number of possible states goes up exponentially. Once the number of sites that are to be considered is determined, the number of possible states for each of those sites and the number of transitions between one state and another needs to be determined when a replication of those sites occur.

When DNA is replicated, it is not a perfect process, occasionally errors occur. The frequency of error can be based on a single replication, and for this analysis, that is how the mutation rate is considered. Upon replication, the base at a given site can be replicated with fidelity giving the same base as the original copy, or the base can be replicated incorrectly and replaced with a different base, a mutation has occurred. Each of those possible transitions on replication has a probability associated with that transition. When considering a single-site DNA evolutionary process, each state will have four possible transitions, and since a single-site DNA evolutionary process has four possible states, it will have 16 (4^2^) possible transitions. When considering a two-site DNA evolutionary process, the 16 possible states will have 256 (16^2^) possible transitions. And when considering the three-site DNA evolutionary process, the 64 possible states will have 4096 (64^2^) possible transitions. Once the number of sites in the genome is determined to be analyzed, the number of states and state transitions can be determined, and the states and transitions can be organized into matrices. The state matrix is a 1 x number of states matrix where the elements of this matrix are the frequencies of each of the possible states of the system at a particular time. The matrix of transitions will be a square dimension equal to the number of possible states where each element of this matrix is the probability of transition from one state to another possible state. These transition probabilities depend on the mutation rate but also depend on the population size. If one assumes that the population size remains constant, the transition matrix will remain constant over time and, a “stationary” transition matrix is obtained and, the transition probabilities remain constant over time. However, if the population size is changing over time, a “non-stationary” transition matrix is obtained and, the transition probabilities will change over time as the population size changes. This is demonstrated by the derivation of the one and two-site DNA evolutionary models for a stationary Markov chain (the Jukes-Cantor model) and the non-stationary variable population model. The evaluation of these Markov chains over time to obtain the frequencies of the different states after each replication is done by simple matrix multiplication.

#### The Markov Process

A Markov chain is a stochastic or random mathematical model describing a sequence of possible events where each event depends only on the state attained in the previous event. Continuous-time Markov chains are called Markov processes and are named after the Russian mathematician Andrey Markov [17]. The relationship between this mathematics and the Kishony Mega-Plate experiment is that with every replication of the bacteria, the offspring potentially get a random mutation that in some cases will allow that variant to make a transition and grow in the next higher drug-concentration region. A well known Markov model of DNA evolution is the Jukes-Cantor Markov Chain model of DNA evolution.[4] There are many derivative models based on the Jukes-Cantor model but none of these models correctly describe the Kishony Mega-Plate experiment. The reason is that the Jukes-Cantor model is based on a stationary transition matrix that gives a Markov process that rapidly reaches equilibrium. The DNA evolutionary process is a highly non-stationary process which does not readily reach equilibrium. An alternative non-stationary transition matrix based on a small modification of the Jukes-Cantor model is presented and the results contrasted with the Jukes-Cantor model.

#### The State Transition Markov Model for a single-site in a Genome

DNA evolution can be mathematically modeled as a discrete Markov Chain. The way this is done for the Kishony Mega-Plate Experiment is as follows. *E*_1_, *E*_2_, *E*_3_, … are the states of the Markov Chain. The way to understand these states for this experiment, consider the first evolutionary step. The wild type bacteria do not have any variants with a mutation which would give them some resistance to the drug these bacteria are challenged. Some site in the bacterial genome lacks the correct DNA base at that site. Replication of a bacterium is the random trial at each transitional step of this Markov process. There are four possible bases for that site, adenine (A), guanine (G), cytosine (C), and thymine (T). Each of those possible bases represents a Markov state. Figure 1 is a diagram of the possible state transitions for this Markov process. The letter μ is the mutation rate that is assumed constant for each of the possible transitions. The state transition diagram for this Markov process is illustrated in Figure 1.

**Illustration 1:**
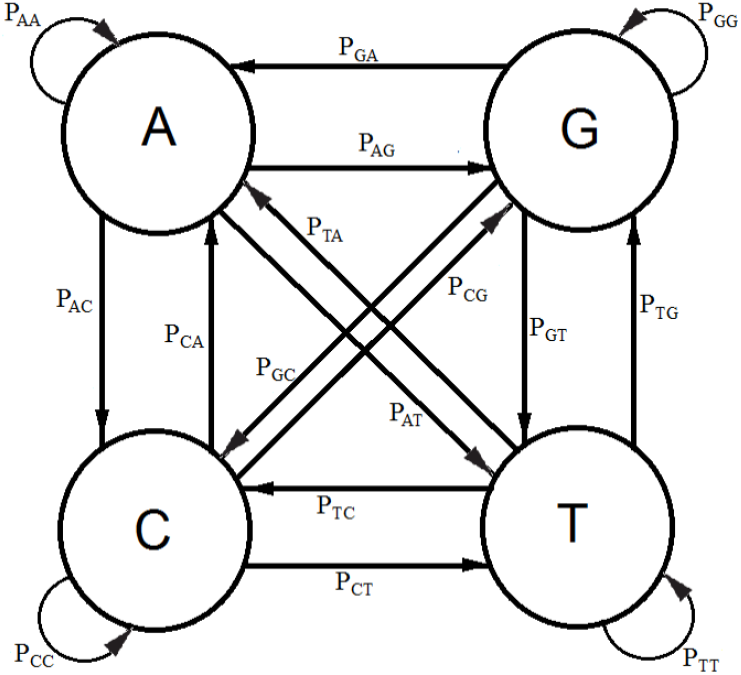
State Transition Diagram for a single mutation at a particular site in the genome.

When replication of the bacterium occurs, if the base at the particular site is T, its descendant can get a T at that site, no mutation occurs, or an A, C, or G base can occur at that site, a mutation occurs. If the base at that site is not a T (that is A, C, or G), then any of the possible bases at that site can be mutated to give a T for that descendant at that particular site. A transition matrix that describes the evolutionary change in time *t* is:

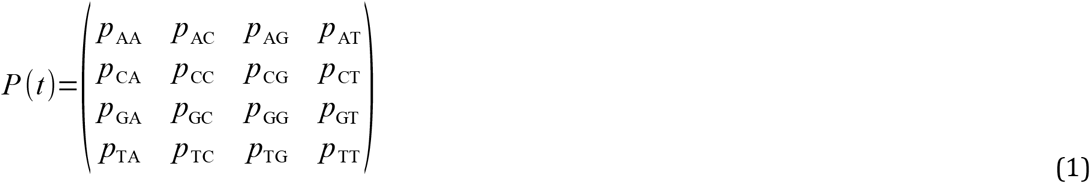

*P* = (*p_ij_*) where the *p_ij_* gives the probabilities of change from the state *E_i_* to *E*_*i*+1_ at time *t* + Δ*t* where Δt is a replication (for the Jukes-Cantor stationary model, it is also a generation). If insertions, deletions, transpositions, and other types of mutations are neglected (that is substitutions only), the transition matrix would look as follows:

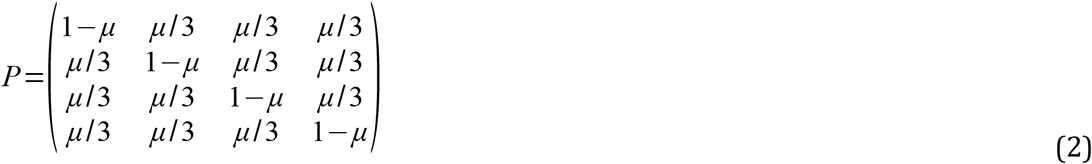

The subscripts on *p* represent each of the possible transitions. For example, an AA subscript means on replication, the adenine base was replicated by another adenine base, no mutation occurs. A CG subscript means that on replication a cytosine base was replaced by a guanine base (a mutation has occurred) and so on. Based on the assumption that the mutation rates are constant, the *p_ij_* elements in terms of the mutation rate can be written as follows and gives the Jukes-Cantor stationary model:

Implicit in the Jukes-Cantor model is the assumption that only a single member of the population is considered. That bacterium replicates, and the Jukes-Cantor model is correct for that first replication. However, for the next replication there are two members of the population, and because of that increase in population size, the change in the relative frequency of the variants must decrease, and that decrease in probabilities for a transition to a different base will not follow the pattern presented by the Jukes-Cantor model. The distribution of variants for the single drug is written as:

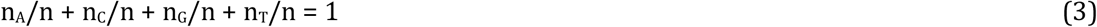

Where n_A_, n_C_, n_G_, and n_T_ are the number of members (replications) with the A, C, G, and T base at the particular site in the population respectively, and n is the total number of members (replications) in the population. For example, if A is assumed to be the variant with the beneficial mutation for the drug and the initial wild-type bacterium has T at that site, then, initially, n_A_, n_C_, and n_G_, are 0, and n_T_ and n are 1. However, with the first replication, there is a small probability that n_A_, n_C_, or n_G_, is 1, a much higher probability that n_T_ is 2, and n will be 2. With each additional replication, there will be a small probability that n_A_, n_C_, or n_G_, will increase by 1, but a much higher probability that n_T_ will increase by 1, and n will increase by 1. With each additional replication, the frequencies of the A, C, and G variants will slowly increase while the frequency of the T variant will very slowly decrease. This effect is not taken into account in the standard Jukes-Cantor stationary model. The probabilities for a variable population model is written:

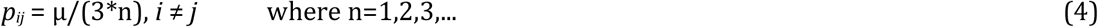

n is the total number of replications (the total population size) in the colony of bacteria. The expected number of members with the mutation A is n_A_=*A*\*n where *A* is the frequency of the A variant after the population has done n replications and so on for the other possible variants. And the corresponding non-stationary Jukes-Cantor transition matrix becomes:

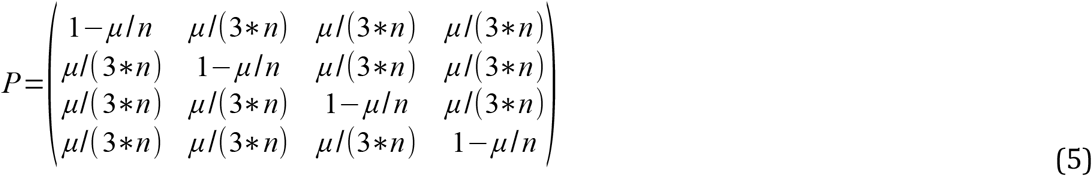

The initial state for either the stationary Jukes-Cantor model or the non-stationary Jukes-Cantor model is written:

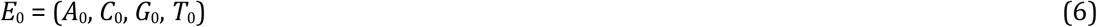

Where the *A*_0_, *C*_0_, *G*_0_, *T*_0_ terms are the frequencies of the particular variant in the initial state. (When A, C, G, or T are italicized indicates the frequencies for the particular variants with that base at that site.) And the state of the system at the time *t_i_* is:

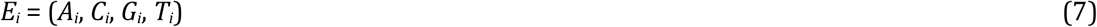

At time *t_i_*, the Markov Chain is in state *E_i_*, then the state of the system at time *t_i_* + Δ*t*, where Δ*t* is one replication, it will be in state *E*_*i*+1_ depends only on *i*, and *t*. The state of the system at time *t*_*i*+1_ is computed by doing the matrix multiplication using the following equation:

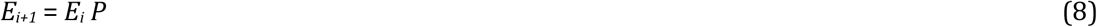

The matrix multiplication for the Jukes-Cantor model is:

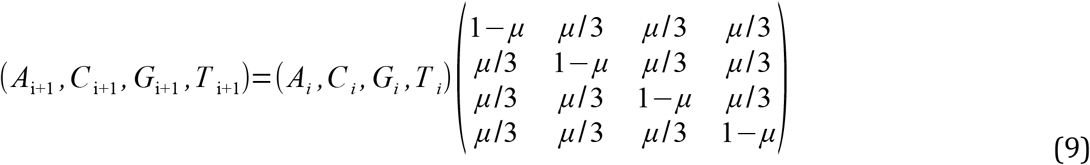

And the matrix multiplication for the non-stationary version of the Jukes-Cantor model is:

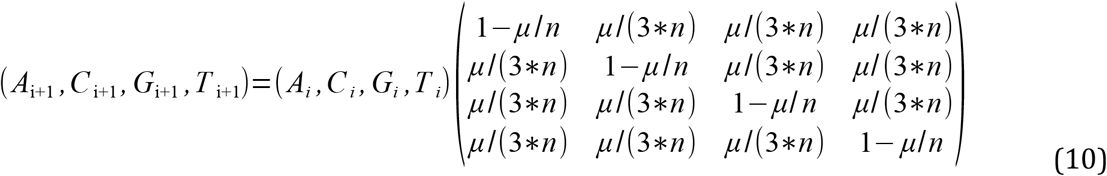

Carrying out the matrix multiplication gives us the four recursion equations which describe this evolutionary Markov process for the stationary and non-stationary models respectively:

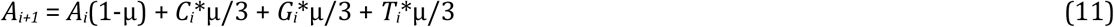

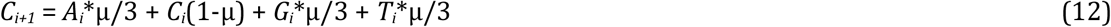

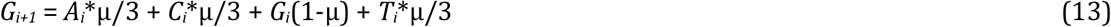

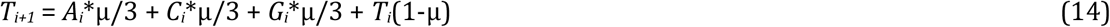

And

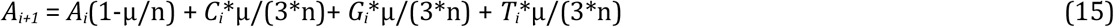

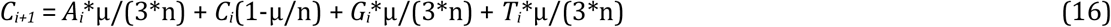

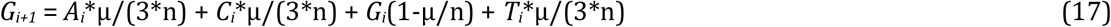

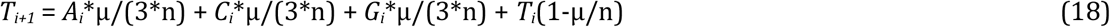

The initial condition for either version using equation (6) is:

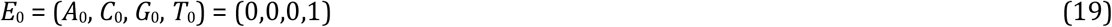

It is assumed that for the Jukes-Cantor model, T is the base at the particular site and for the Kishony Mega-Plate experiment that the initial inoculate at time = 0, the bacterium has a T base where an A base is needed to give resistance. The fundamental difference between the stationary and non-stationary models is that as the population size grows, n is increasing in the non-stationary model. The stationary process assumes that the population size is constant (n=1) for all time.

FORTRAN computer programs were written to evaluate both stationary and non-stationary versions and compared for two different mutation rates, μ = 1E-5 and μ = 1E-9 and, are presented in the ***Results*** section. An equivalent calculation was carried out based on two-site models for stationary and non-stationary Markov processes.

#### The Markov Process for Two Sites, the Jukes-Cantor stationary model and the Modified Jukes-Cantor non-stationary model

The analysis for the DNA evolutionary process for two sites begins with the construction of the state transition diagram for the evolution at two sites (Figure 2). This diagram is much more complex in that there are 16 possible states. (Note that only the transition lines to and from state A1A2 to the other states are included in this figure. Transition lines from the other states would appear similarly). In Figure 2, the possible state transitions are drawn for a member of the population which has an A base at site 1 and an A base at site 2 and the possible transitions that can occur on replication. An A1A2 member can replicate and produce another A1A2 member, or an A1C2, or a C1A2, or any of 13 other possible transitions.

**Illustration 2:**
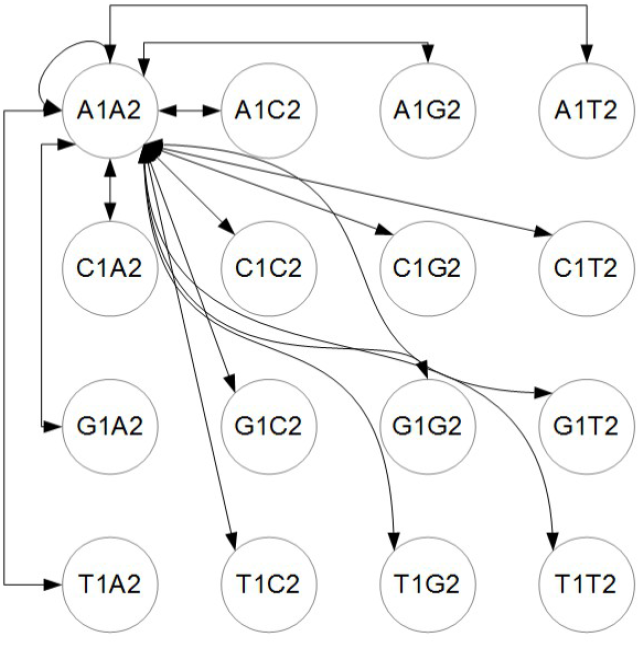
The two-site Transition Diagram, only A1A2 transition lines shown, similar transition lines for other states apply but not shown.

The illustration in Figure 3 gives the diagram for a single state with the understanding that equivalent states exist for the other possible combinations of bases. The number following the base letter indicates the site.

**Illustration 3:**
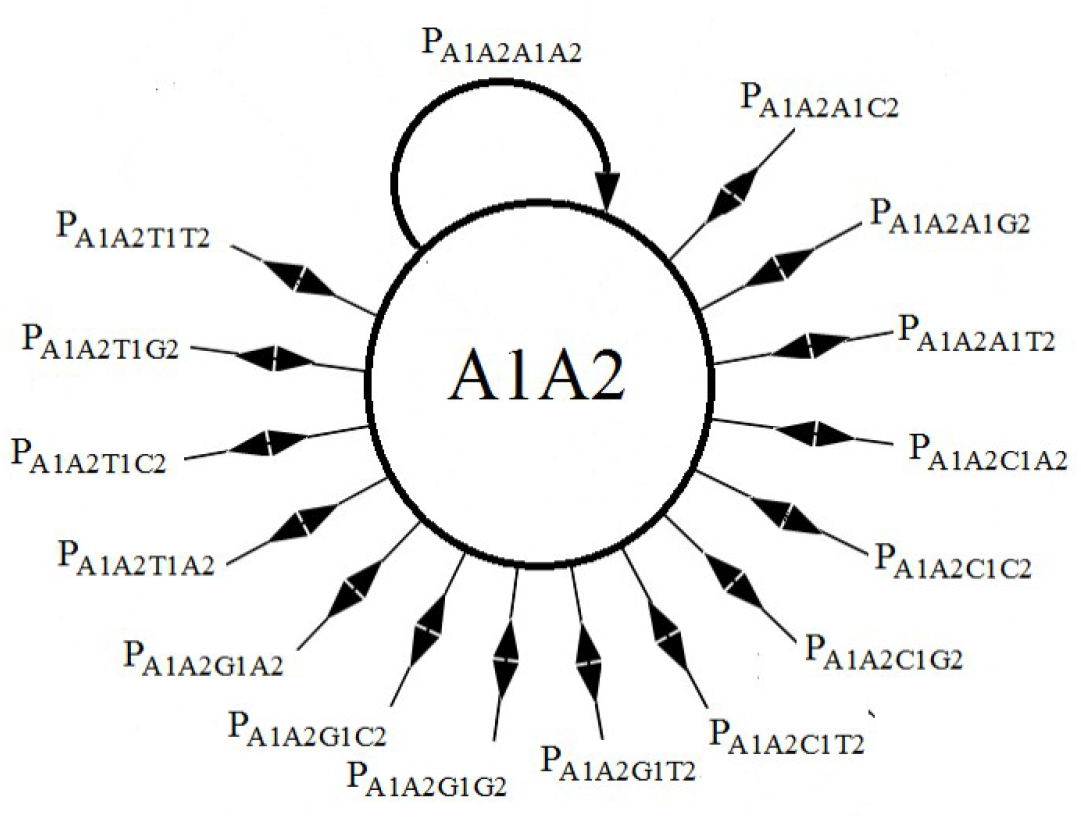
Possible state transitions for two sites from an A at site 1 and an A at site 2 to all other possible transitional states.

In contrast with the single-site model which has 4 possible states and 4 possible transitions from each state, the two-site (second-order) model has 16 possible states with 16 possible transitions from each state to the other possible states. The list of possible transitions for the A state for the single-site model gives 4 probabilities, *p*_AA_, *p*_AC_, *p*_AG_, and *p*_AT_. The transition probabilities for the A1A2 state for the two-site model is, *p*_A1A2A1A2_, *p*_A1A2A1C2_, *p*_A1A2A1G2_, *P*_A1A2A1T2_, *p*_A1A2C1A2_, *p*_A1A2C1C2_, *P*_A1A2C1G2_, *p*_A1A2C1T2_, *p*_A1A2G1A2_, *p*_A1A2G1C2_, *p*_A1A2G1G2_, *p*_A1A2G1T2_, *p*_A1A2T1A2_, *p*_A1A2T1C2_, *p*_A1A2T1G2_, and *p*_A1A2T1T2_. Similar probabilities are obtained for the other 240 possible state transitions in the two-site model. A *p*_A1A2A1A2_ transition means that an A base is at site 1 and an A base is at site 2 in the parent, and that on replication, the descendant will also have an A base at site 1 and an A base at site 2, no mutation has occurred on replication at either site. A *p*_A1A2A1C2_ transition means that an A base is at site 1 and an A base is at site 2 in the parent and that on replication, the descendant will also have an A base at site 1 but a C base at site 2, no mutation has occurred on replication at site 1 but a C mutation has occurred at site 2. A *p*_A1A2C1G2_, transition means that an A base is at site 1 and an A base is at site 2 in the parent and that on replication, the descendant will have a C base at site 1 and a G base at site 2, a C mutation has occurred on replication at site 1 and a G mutation has occurred at site 2. And so on for the other possible transitions.

For the evolution of two sites, a 16×16 transition matrix is obtained which is partially displayed in equation (20).

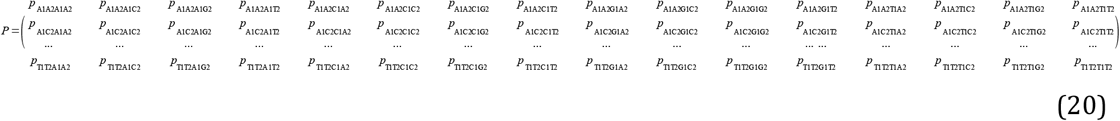

If n is assumed constant and equal to 1, the first row of the two-site transition matrix in terms of the mutation rate give the Jukes-Cantor stationary version of the two-site model:

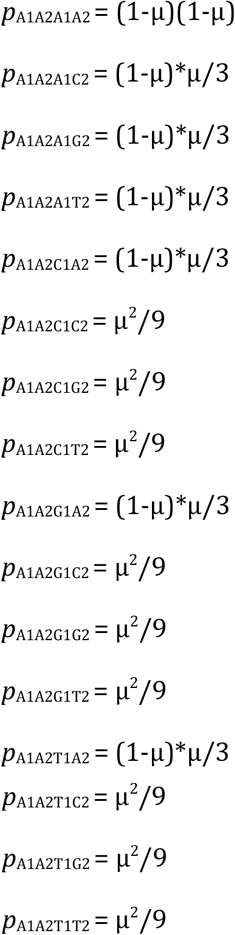

The other rows of the transition matrix from the 240 other possible state transitions will have similar probabilities. These values are substituted into the two-site transition matrix for the Jukes-Cantor two-site stationary model. This 16×16 matrix is partially displayed in equation (21).

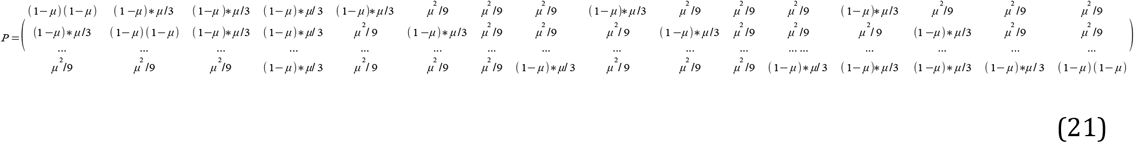

The Jukes-Cantor two-site non-stationary model has the following transition probabilities based on the same reasoning for the single-site model. The probabilities for the A1A2 state are written as:

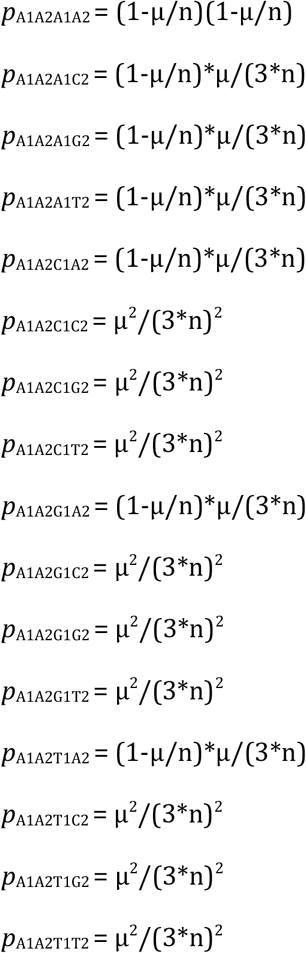

The other rows of the non-stationary transition matrix from the 240 other possible state transitions have similar probabilities. These values are substituted into the two-site transition matrix for the Jukes-Cantor two-site non-stationary model. This 16×16 matrix is partially displayed in equation (22).

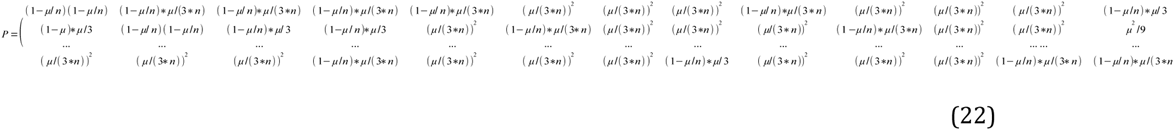

In a similar manner as with the single-site model, the state transition equation is:

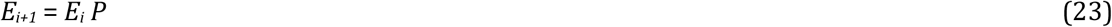

Equations (21) and (22) are used for the stationary and non-stationary models (and the 15 other equivalent transition equations) and the two-site transition matrix to compute the Markov chain recursion equations.

(Note that when X1Y2 where X and Y can be A, C, G, or T are italicized indicates the frequencies of those variants with X at site 1 and Y at site 2.) The recursion equation for the A1A2 state Jukes-Cantor stationary model is:

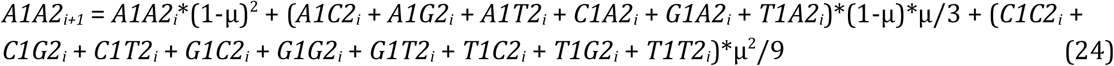

Equivalent equations are obtained for the other 15 states. The equivalent recursion equation for the A1A2 state Jukes-Cantor non-stationary model is:

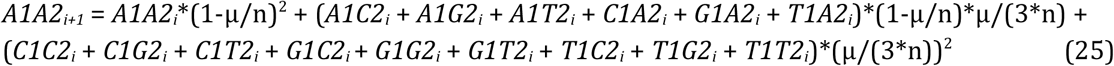

Similar equations for the other 15 possible states give the full definition of the Markov transformation for the non-stationary model.

The initial state for either two-site system is written:

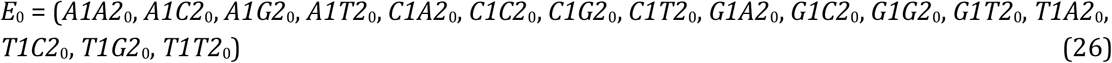

The *A1A2*_0_, …, *T1T2*_0_ terms are the frequencies of the particular variant in the initial state. The state of the system at time *t*_*i*+1_ is given by equation (23).

It is assumed that in the initial state, the wild type variant inoculated on the plate for a two-drug experiment has bases T at site 1 and at site 2 such that T1T2_0_ = 1 (that is the frequency of that variant is 1) and the other 15 possible states *A1A2*_0_, …, *T1G2*_0_ = 0 gives an initial condition of:

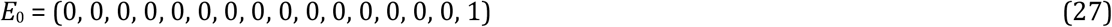

The stationary and non-stationary two-site models were evaluated similarly as the single-site models and compared for two different mutation rates, μ = 1E-5 and μ = 1E-9, and presented in the **Results** section.

## Results

The Numerical Calculation of the Jukes-Cantor single-site Stationary and Non-stationary

### Models

FORTRAN computer programs were written to compute the values for equations (11–14) stationary model and equations (15–18) non-stationary model for two mutation rates, 1E-5 and 1E-9. The FORTRAN source code is supplied in the supplemental documentation as well as the data derived from these programs. The results are shown in the graphs below for the Jukes-Cantor stationary and non-stationary models for two mutation rates. The first two figures are for the single site stationary systems (mutation rates 1E-5 and 1E-9 respectively.

The base frequencies for the C and G variants are not included in Figure 4 because these base frequencies are identical to the A frequency curve. This is because of the symmetry of the Jukes-Cantor model. The mutation rate for base transitions is the same for base transversions. Each Markov transition in the Jukes-Cantor stationary model is both a replication and a generation. Each element in the transition matrix is the mutation rate for a particular mutational change in a single replication for a single member of a lineage. Therefore, each replication is also a generation.

**Illustration 4:**
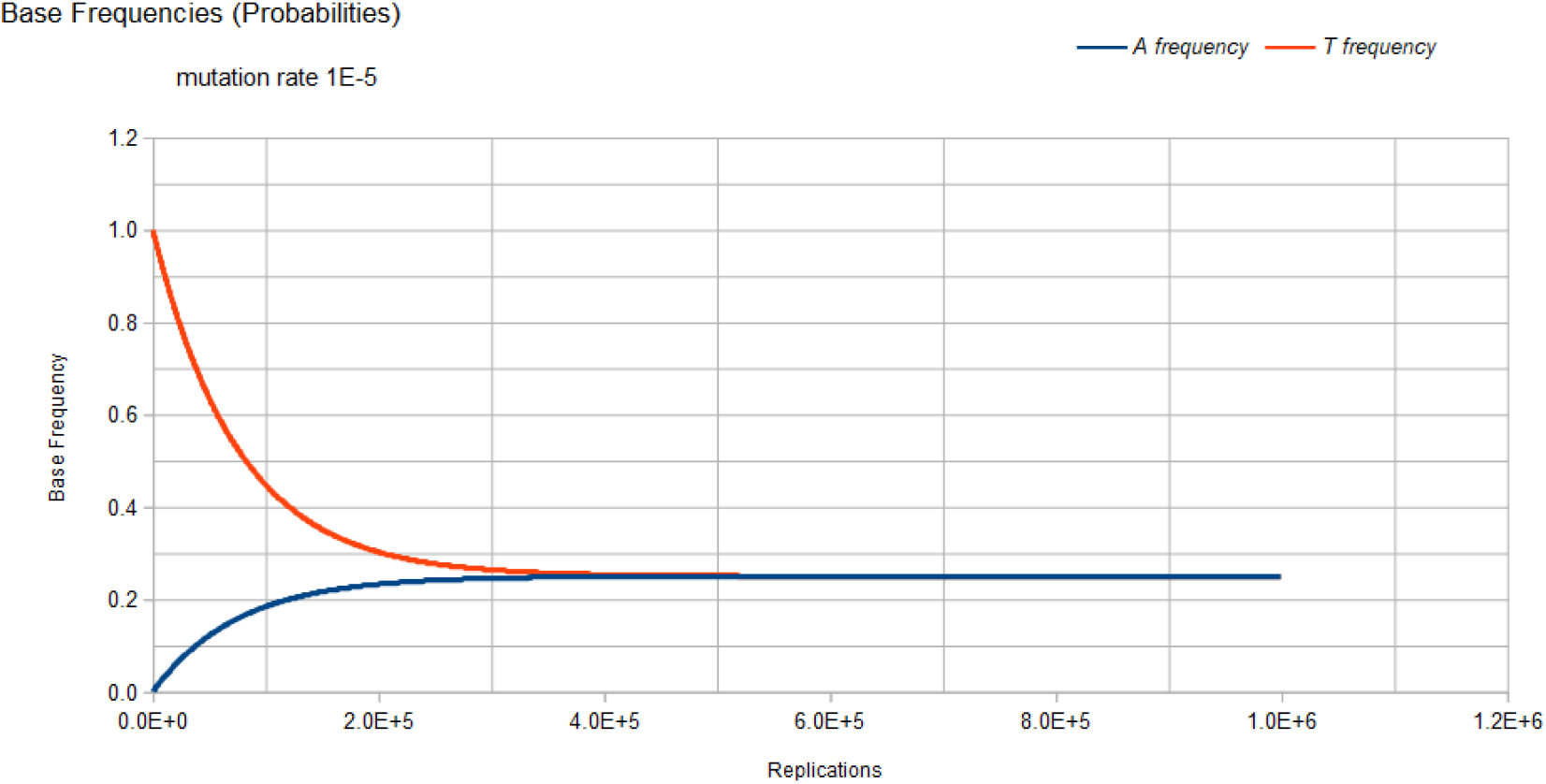
Base frequencies for A and T variants, Jukes-Cantor stationary model, single-site, as a function of number of replications and mutation rate 1E-5

The way to correlate the curves in Figure 4 to the genetic transformation is to consider some founder bacterium of a lineage in the population with a T base at a particular site in the genome. That bacterium replicates, and its descendant will have a slightly reduced probability of a T base at that site and a slightly increased probability of having an A, C, or G base (mutation) at that site. When that descendant replicates, its descendant will have another slight decrease in the probability of a T base occurring at that site and another slight increase in the probability of having an A, C, or G mutation at that site. Each replication of the following descendants will decrease the probability of T occurring at that site and increase the probability of an A, C, or G mutation at that site until the evolutionary process reaches equilibrium where the probability of any of the four possible bases occurring at that site will be 0.25. For the mutation rate of 1E-5, this occurs at about 4E5 replications (generations). After 4E5 replications, the probability of finding any of the four bases at that site remains constant at the value of 0.25. This evolutionary model has reached equilibrium.

Figure 5 demonstrates the result of the Jukes-Cantor stationary model similar to Figure 4 except with a lower mutation rate. As with Figure 4, Figure 5 does not include the C and G frequency (probability) curves because these base frequencies are identical to the A frequency curve. The same Markov DNA evolutionary process described in Figure 4 is occurring with the case whose result is illustrated in Figure 5. The only difference is that the lower mutation rate (1E-9 vs. 1E-5) results in a slower approach to equilibrium (4E9 vs. 4E5 replications), but either case converges on the same equilibrium values for the base frequencies, 0.25.

**Illustration 5:**
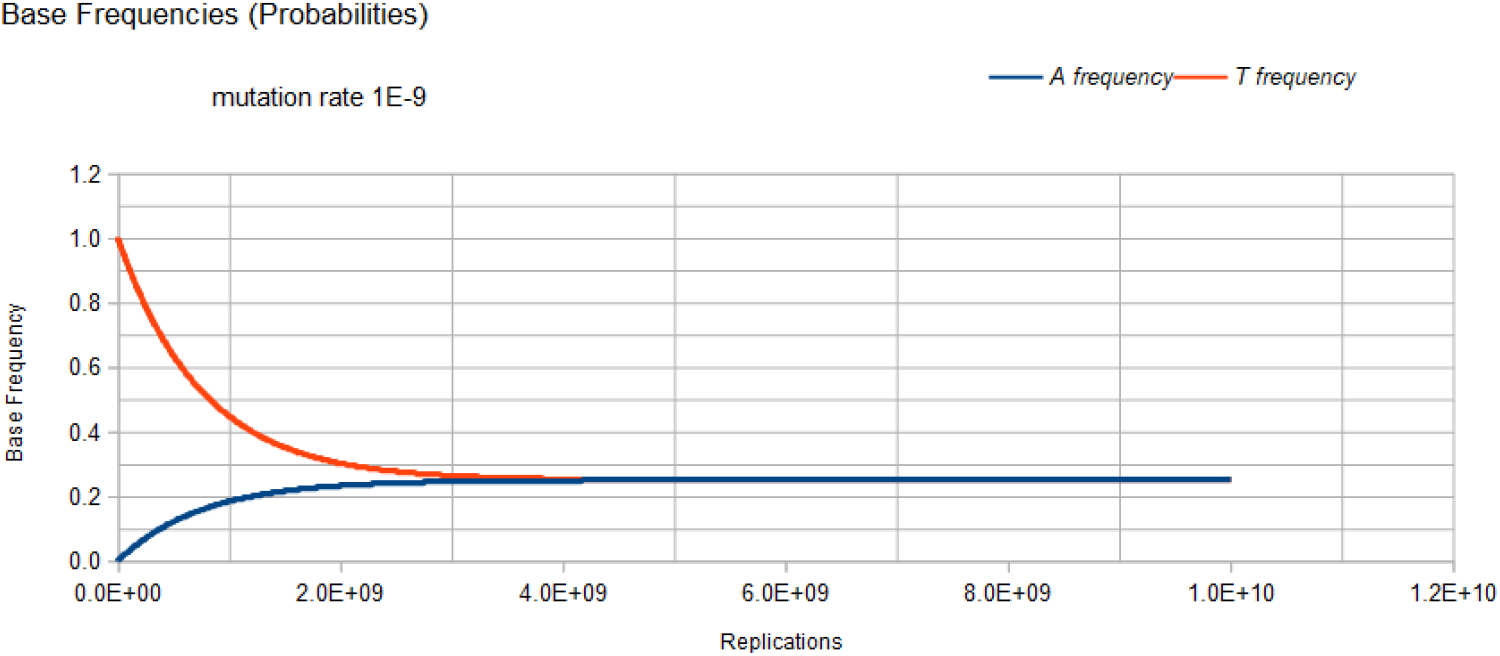
Base frequencies for A and T variants, Jukes-Cantor stationary model, single-site, as a function of number of replications and mutation rate 1E-9

Figure 6 gives the result of the Jukes-Cantor non-stationary model of DNA evolution. It is this model that is proposed to be the correct simulation of the Kishony Mega-Plate experiment. As with the Jukes-Cantor stationary model, as described in Figures 4 and 5, the frequencies (probabilities) for the C and G variants are not plotted because these base frequencies give identical curves to the A frequency curve. In this case, each replication is not a generation. To understand this graph (and mathematics) in the context of the Kishony Mega-Plate experiment, consider what happens from the start of the initial condition of the experiment.

**Illustration 6:**
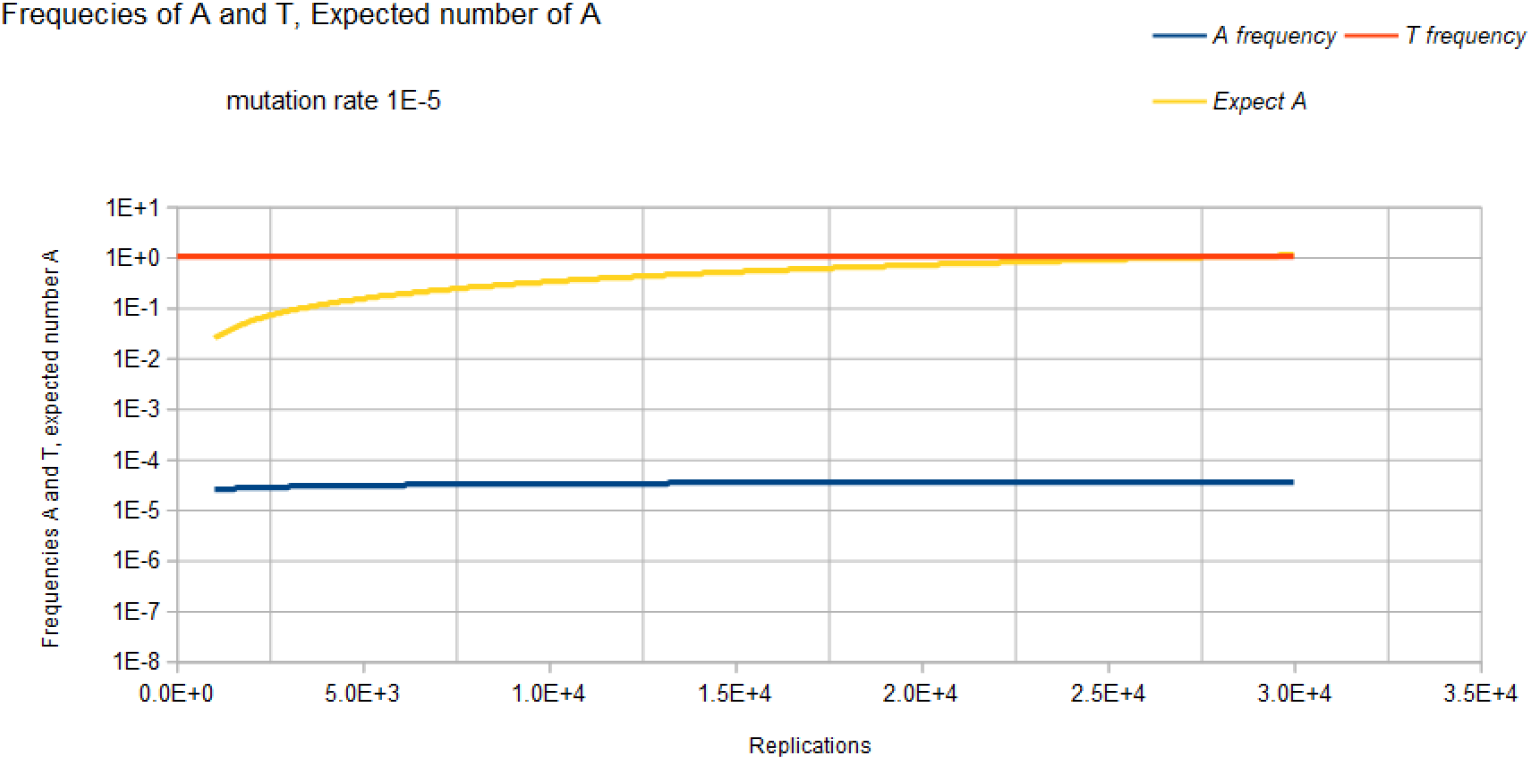
Base frequencies for A and T variants and expected number of A, Jukes-Cantor non-stationary model, single-site, as a function of number of replications and mutation rate 1E-5.

Initially, a non-drug resistant bacterium is inoculated into the drug-free region of the petri dish. In this case, it is assumed that the base at the site of interest is T. Initially, the frequency of that variant is 1, and the frequency of any variant with an A, C, or G base at that site is 0. That bacterium double for the first generation. The Markov transition matrix for that single replication gives a small probability that an A, C, or G mutation occurs, and the probability that a T occurs is slightly less than 1. These two bacteria double for the second generation. It requires 2 Markov transitional steps to compute the frequencies of the different variants for this generation. Each of these Markov transitional steps slightly reduces the frequency of the T variants and slightly increases the frequencies of the A, C, and G variants. The next generation consists of a doubling of these four bacteria to 8 bacteria that requires 4 Markov transitional steps to compute the frequencies of each of the different variants for this generation. This DNA evolutionary process continues until the frequency of A, and the total population size is sufficient to give an expected occurrence of an A variant (n_A_=n\**A*). For a mutation rate of 1E-5, this occurs at about a population size of 3E4 (about 15 doublings or generations).

Another consideration is that this Markov process is occurring at every site in the genome of the bacteria used by the Kishony team. Two different antimicrobial agents were used (not simultaneously) in the experiment, Ciprofloxacin, and Trimethoprim. The occurrence of resistance mutations to either drug occurs in a single colony which is demonstrated by the use of either drug in the experiment. The experiment does not work when both drugs are used simultaneously [18]. In order for the experiment to work with 2 drugs requires that a single variant have resistance mutations for both drugs. The mathematical requirement for this to occur is demonstrated by the 2 sites non-stationary Markov process.

Figure 7 demonstrates the results of the Jukes-Cantor non-stationary model similar to Figure 6 except with a lower mutation rate. As with Figure 6, Figure 7 does not include the C and G frequency (probability) curves because these base frequencies are identical to the A frequency curve. The same Markov DNA evolutionary process that is described in Figure 6 is occurring with this case whose result is illustrated in Figure 7. The only difference is that the lower mutation rate (1E-9 vs. 1E-5) results in a slower approach to an expected occurrence of variant A equal to 1. (1.5E8 (about 28 doublings or generations) vs. 3E4 replications (about 15 doublings or generations)).

**Illustration 7:**
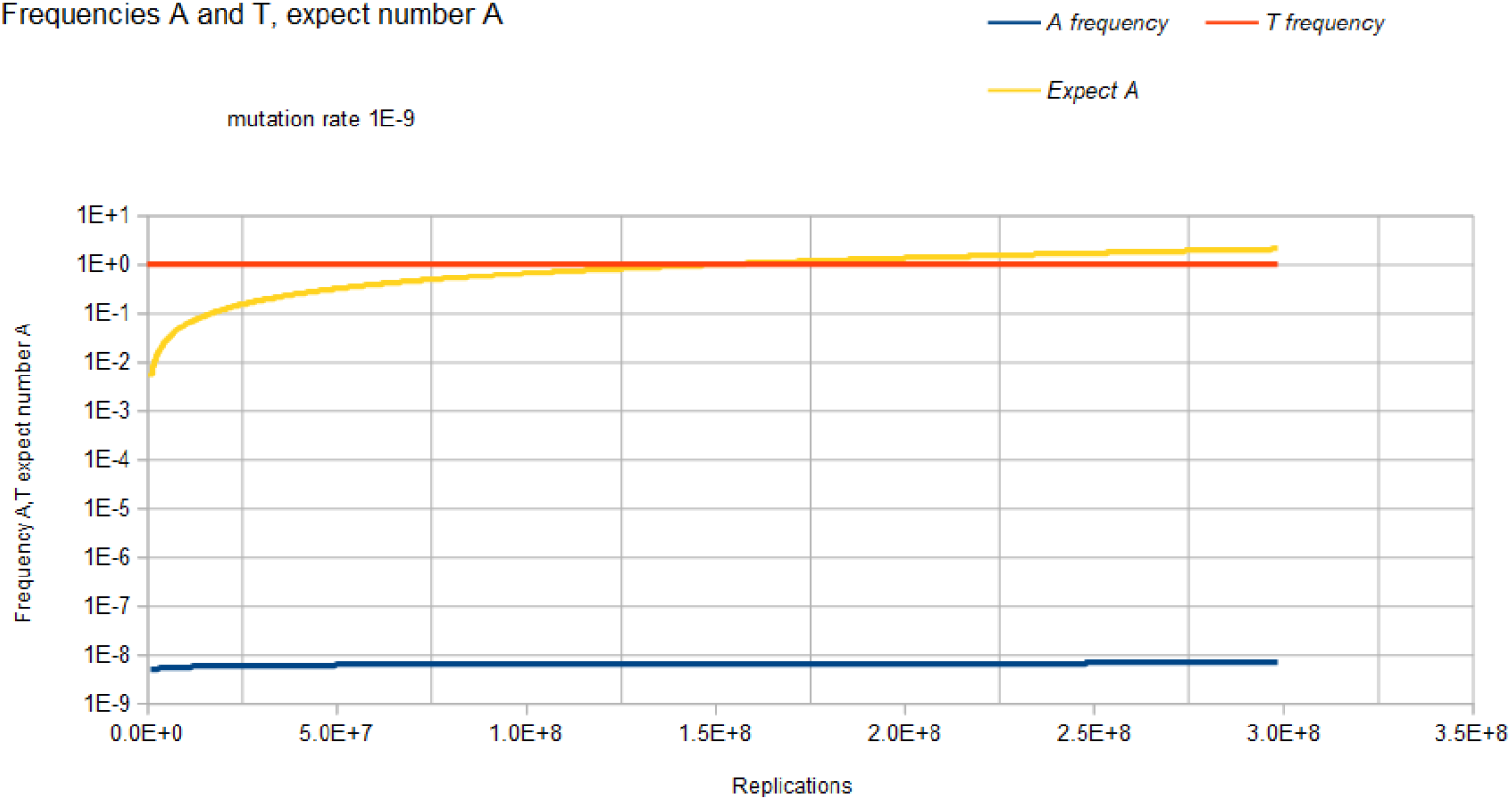
Base frequencies for A and T and expected number of variant A, Jukes-Cantor non-stationary model, single-site, mutation rate 1E-9.

The following four figures show the results for the two site Jukes-Cantor model, the first two of these four are for the stationary two sites model, mutation rates 1E-5 and 1E-9 respectively, and the last two figures for the two sites non-stationary model, mutation rates 1E-5 and 1E-9 respectively.

FORTRAN computer programs were written to compute the values for equation (25) (plus the 15 other equivalent equations) stationary model and equation (26) (plus the 15 other equivalent equations) non-stationary model for two mutation rates, 1E-5, and 1E-9. The results are shown in the following two graphs below for the two sites Jukes-Cantor model (mutation rate 1E-5 and 1E-9 respectively). Source code and computed data are included in supplementary documentation.

The base frequencies for the X1Y2 variants where neither X nor Y is a T base are not included in Figure 8 because these base frequencies are identical to the A1A2 frequency curve. The base frequencies for the X1T2 and T1Y2 variants where neither X nor Y is a T base are not included in Figure 8 because these base frequencies are identical to the A1T2 frequency curve. This is again because of the symmetry of the Jukes-Cantor model. The mutation rate for base transitions is the same for base transversions. Each Markov transition in the Jukes-Cantor stationary model is both a replication and a generation. Each element in the transition matrix is the product of the individual mutation rates (that is the joint probability) of the particular mutational change at each site in a single replication for a single member of a lineage. Therefore, each replication is also a generation.

**Illustration 8:**
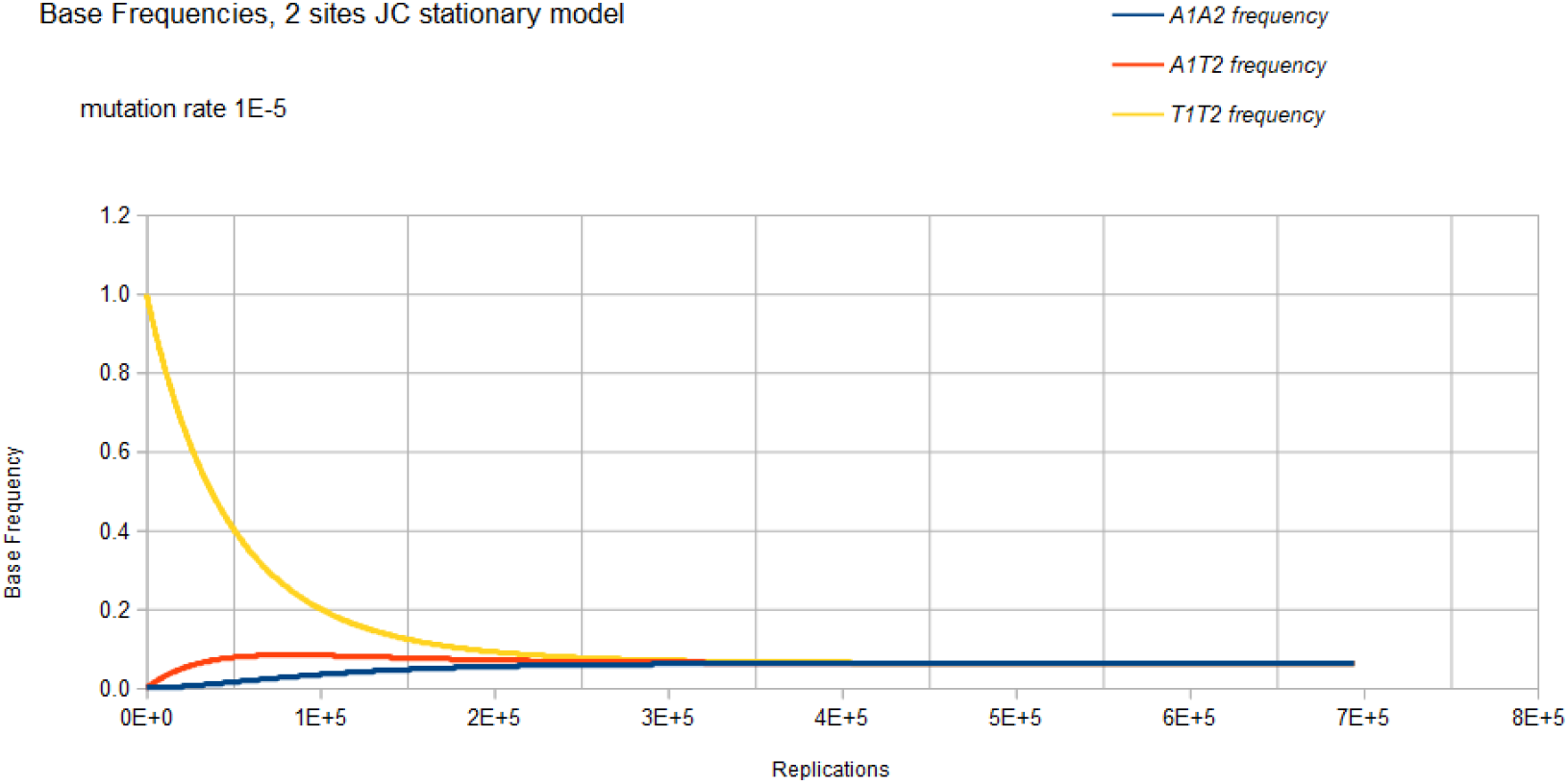
Base frequencies for A1A2, A1T2, and T1T2 variants, Jukes-Cantor stationary model, two sites, as a function of number of replications and mutation rate 1E-5

The way to correlate the curves in Figure 8 to the genetic transformation is to consider a bacterium in a population with a T1 base at one particular site and a T2 base at another particular site in the genome. That bacterium replicates, and its descendant will have a slightly reduced probability of either a T1 base at the one particular site or a T2 base at the other particular site and a slightly increased probability of having an A1, A2, C1, C2, G1, or G2 base (mutation) at their particular sites. When that descendant replicates, its descendant will have another slight decrease in the probability of a T1 or T2 base occurring at their particular sites and another slight increase in the probability of having an A1, A2, C1, C2, G1, or G2 mutation at their particular sites. Each replication of each of the following descendants will decrease the probability of T1 or T2 occurring at the particular sites and increase the probability of an A1, A2, C1, C2, G1, or G2 mutation at their particular sites until the evolutionary process reaches equilibrium where the probability of any of the eight possible bases (four possible bases at each site) occurring at that site will be 0.0625. For the mutation rate of 1E-5, this occurs at about 4E5 replications (generations). After 4E5 replications, the probability of finding any of the eight possible bases at the two particular sites remains constant at the value of 0.0625. This evolutionary model has reached equilibrium.

Figure 9 demonstrates the result of the Jukes-Cantor two-site stationary model similar to Figure 8 except with a lower mutation rate. As with Figure 8, Figure 9 does not include most of the base frequencies for the same reason, as described in Figure 8. The only difference is that the lower mutation rate (1E-9 vs. 1E-5) results in a slower approach to equilibrium (4E9 vs. 4E5 replications (generations)), but either two sites stationary case converges on the same equilibrium values for the base frequencies, 0.0625.

**Illustration 9:**
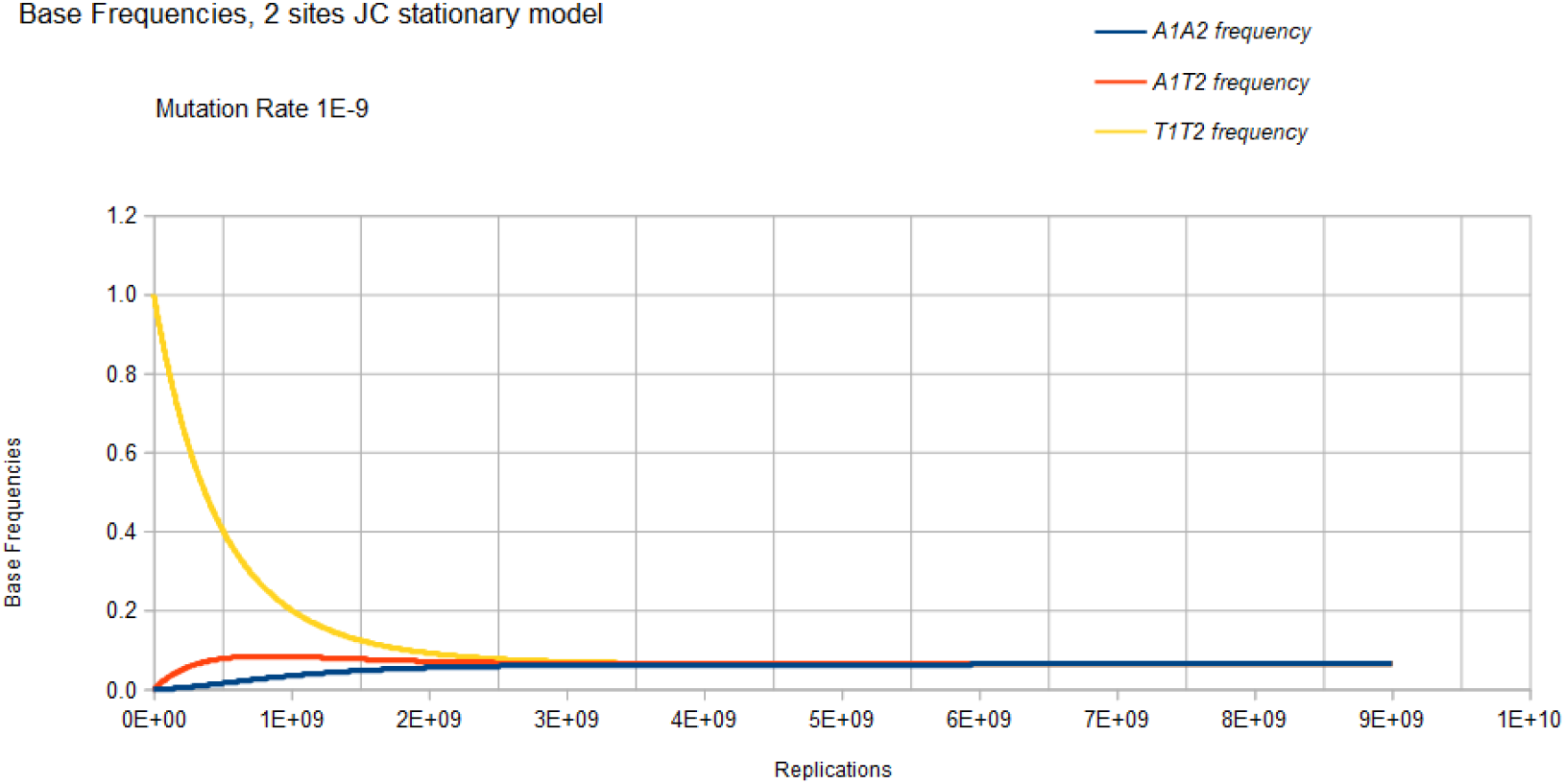
Base frequencies for A1A2, A1T2, and T1T2 variants, Jukes-Cantor stationary model, two sites, as a function of number of replications and mutation rate 1E-9

Figure 10 gives the result of the Jukes-Cantor two-site non-stationary model of DNA evolution. The base frequencies for the X1Y2 variants where neither X nor Y is a T base are not included in Figure 10 because these base frequencies are identical to the A1A2 frequency curve. The base frequencies for the X1T2 and T1Y2 variants where neither X nor Y is a T base are not included in Figure 10 because these base frequencies are identical to the A1T2 frequency curve. This is again because of the symmetry of the Jukes-Cantor model. The mutation rate for base transitions is the same for base transversions. Each Markov transition in the Jukes-Cantor 2 site non-stationary model is only a replication, not a generation. Each element in the transition matrix is the product of the individual mutation rates (that is the joint probability) of the particular mutational change at each site in a single replication where the change in frequencies of the different variants now depends on population size.

**Illustration 10:**
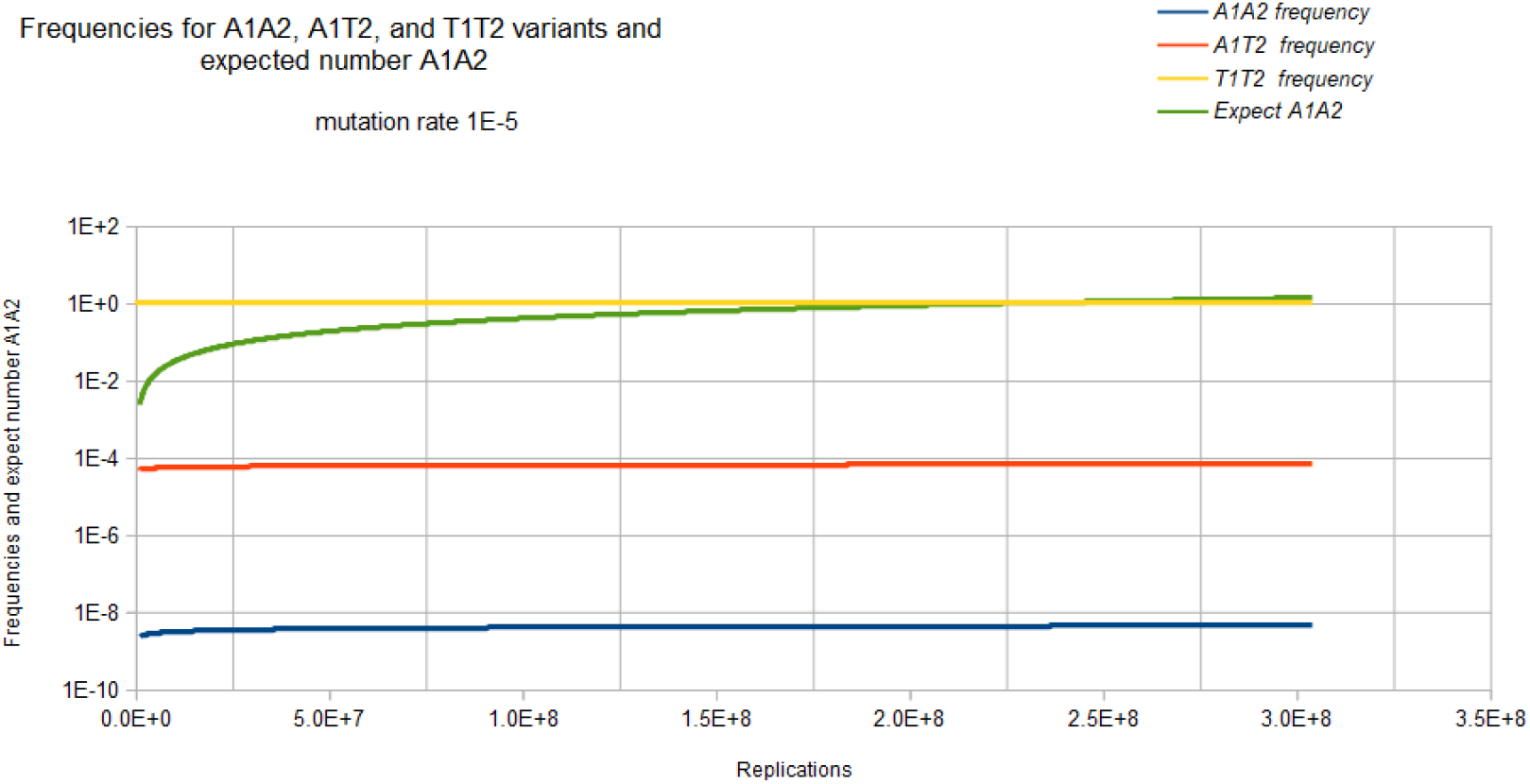
Base frequencies for A1A2, A1T2, and T1T2 variants, and expected number of A1A2 variants, Jukes-Cantor non-stationary model, two sites, as a function of number of replications and mutation rate 1E-5.

Consider the math presented here in the context if the Kishony team tries to perform the experiment with two drugs (or if the step increase in drug concentration is so large that two mutations are required for adaptation to the next higher drug concentration region). Initially, a single non-drug resistant bacterium is inoculated into the drug-free region of the petri dish. In this case, it is assumed that the base at one site of interest is T1 and for the second site of interest the base is T2 so that the frequency of that T1T2 variant is 1 and the frequency of any variant with combinations A1, A2, C1, C2, G1 or G2 base at sites 1 and 2 are 0. That bacterium double for the first generation. The Markov transition matrix for that single replication gives a small probability that an A, C, or G mutation occurs at either site and the probability that a T occurs at either site is slightly less than 1. These two bacteria double for the second generation. This requires 2 Markov transitional steps to compute the frequencies of the different possible variants. Each of these Markov transitional steps slightly reduce the frequency of the T1T2 variant and very slightly increase the frequencies of the A1A2, A1C2, A1G2, C1A2, C1C2, C1G3, G1A2, G1C2, and G1G2 variants and slightly increase the frequencies of the A1T2, C1T2, G1T2,T1A2, T1C2, and T1G2 variants. The next generation consists of a doubling of these four bacteria to eight bacteria which requires 4 Markov transitional steps to compute the frequencies of each of the different variants. This DNA evolutionary process continues until the frequency of A1A2 (the assumed double drug resistant variant) and the total population size is sufficient to give an expected occurrence of an A1A2 variant (nA1A2=n*A1A2). For a mutation rate of 1E-5 this occurs at about a population size of 2.3E8 (about 28 doublings or generations).

Figure 11 gives the result of the Jukes-Cantor 2 site non-stationary model similar to Figure 10 except with a lower mutation rate. As with Figure 10, Figure 11 does not include A1C2, A1G2, C1A2, C1C2, C1G2, G1A2, G1C2, and G1G2 frequency curves because these base frequencies are identical with the A1A2 frequency curve. The base frequencies for the C1T2, G1T2, T1A2, T1C2, and T1G2 variants are not included in Figure 11 because these base frequencies are identical to the A1T2 frequency curve. The frequency of the T1T2 variant remains very close to 1, only very slowly decreasing as the number of replications increases so it is not displayed. The A1A2 frequency curve appears superimposed on the A1T2 curve because of the very small values (of the order of 1E-16 and 1E-8 respectively). The expected occurrence of an A1A2 variant is very slowly increasing and the computation was halted at 3E14 replications (about 48 doublings (generations)). A linear interpolation of the data from Figure 11 gives the expected occurrence of an A1A2 variant will occur at about 8E15 replications (about 53 doublings of the original founder bacterium). For the Kishony Mega-Plate experiment to work with two drugs (or a larger step increase in drug-concentration), a much, much larger petri dish will be needed than the “Mega-Plate”.

**Illustration 11:**
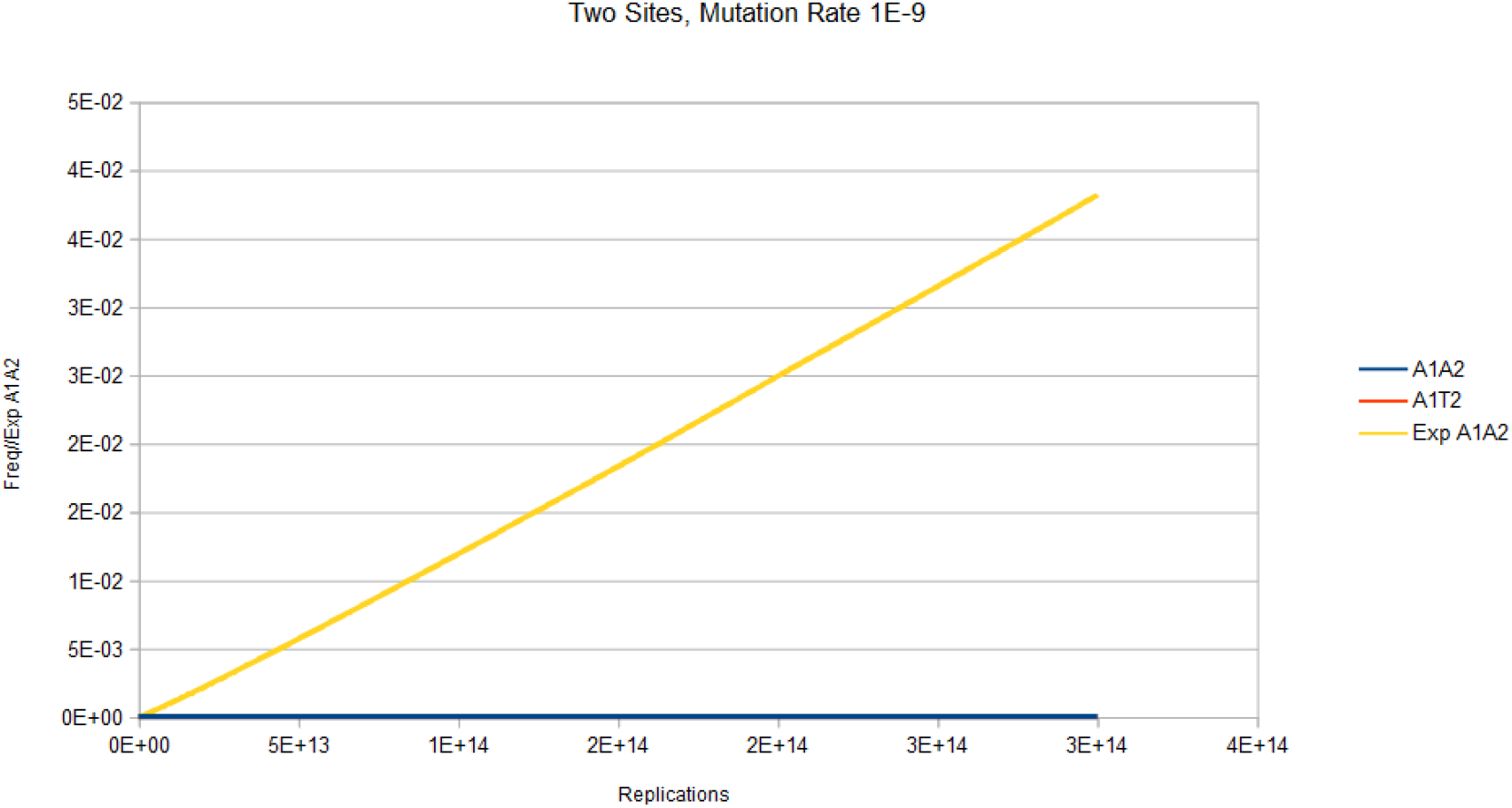
Base frequencies for A1A2 and A1T2 variants, and expected number of A1A2 variant, Jukes-Cantor non-stationary model, two sites, as a function of number of replications and mutation rate 1E-9

## Discussion/Conclusion

The mathematical behavior of the Markov process of DNA evolution is significantly different when assuming a stationary versus a non-stationary model. It is shown that the Jukes-Cantor model assumes a constant population size of 1 because of the elements used in the transition matrix. The Jukes-Cantor stationary model is physically modeling a single-site in the DNA in a lineage of a single member of a population. Because of this, each transition from one state to the next represents a generation. The initial state at that site, the probability (frequency) of that base is 1, and the probability of any of the other bases is 0. When that member replicates, the probability of the original base occurring is slightly less than 1 and the probability of any of the other substitutions now is slightly greater than 0. When that descendant replicates, the probability of the original base is again slightly decreased and the probabilities of any of the possible substitutions are increased. When the equilibrium state is achieved, the probabilities of finding any of the possible bases will be 0.25 for all possible bases. And the number of replications to reach that state is approximately 1/(¼*μ). Once the equilibrium state is reached, with any further replications, the probability of finding any base at that site will be 0.25 no matter how many further replications occur.

The Jukes-Cantor model for a single-site converges to a stationary value of 0.25 regardless of the mutation rate, and the two-site model converges to 0.0625 (again regardless of the mutation rate). As with, the single-site Jukes-Cantor model, the equivalent two-site model also applies to the lineage of a single member where each transition represents a replication of that particular member, but that replication also represents a generation. When a constant non-unity population is considered with the model, each matrix multiplication still represents a single replication of that member, but now a generation is n matrix multiplications (replications). The non-stationary Jukes-Cantor model takes into account the number of replications (population size) occurring in a lineage. This has a strong effect on the relative frequencies of the different variants in the population. This model does not reach equilibrium. The Kishony Mega-Plate experiment starts its evolutionary process with a single “wild-type” variant. As these wild-type replicates and the population size increases, the relative frequencies are changing, but very slowly, and most of the members of the population will have the original base at the site of interest. The probability of the correct mutation occurring is improving as the colony size grows, but its relative frequency will be very low because of the large population size, and this is demonstrated by the non-stationary Jukes-Cantor model as well as the Kishony Mega-Plate Experiment.

The single-site Jukes-Cantor stationary model with a mutation rate of 1E-5 reaches equilibrium and relative frequencies of all variants of 0.25 at about 4E5 replications, as shown in Figure 4. For the same mutation rate and the number of replications, the single-site Jukes-Cantor non-stationary model shows the relative frequency of the wild-type to be very close to 1, and the relative frequency of the mutated variant to be 6E-5, but the expected number of the drug-resistant mutated variant will be about 1. Likewise, for a mutation rate of 1E-9, the single-site Jukes-Cantor stationary model reaches an equilibrium of 0.25 at about 3E9 replications. This is the same number of replications that on average would give every possible base substitution at every site in a genome. And this is the approximate number of replications for the Kishony populations to get the next beneficial mutation for the next higher drug-concentration region.

The reason why such significant differences occur between the stationary and non-stationary models can be seen when equations (4) and (16) are considered, *E_i_* = (*A_i_, C_i_*, *G_i_, T_i_*) for the single-site model that is identical for both stationary and non-stationary models. *E_i_* is the state of the system at time (generation), *i* and *A_i_, C_i_, G_i_, T_i_* are the frequencies of the bases in that population at time *i*. If the total population size is n, and the number of members of the population of each of the variants is n_Ai_, n_Ci_, n_Gi_, n_Ti_ respectively, then frequencies *A_i_, C_i_, G_i_, T_i_* at time *i* will be n_Ai_/n, n_Ci_/n, n_Gi_/n, n_Ti_/n, respectively. In the stationary formulation of the Jukes-Cantor model, n is implicitly assumed to be constant, and equal to 1. In the Kishony Mega-Plate experiment, n is not constant and varies with time. When n is considered to be a function of time as demonstrated above, the non-stationary Jukes-Cantor model will give accurate predictions of this experiment. This principle becomes even more apparent for Markov Chain models of DNA evolution when two sites are being considered simultaneously.

The two-site stationary model with a mutation rate of 1E-9 also reaches equilibrium at about 3E9 replications and a frequency value of 0.0625. This value under-predicts the number of replications in the Kishony experiment to accumulate the first two beneficial mutations sequentially by a factor of 2. Also, the frequency of the different variants will be nowhere close to 0.0625. In the actual experiment, the frequency of the wild-type is still very close to 1, and the mutated members of the population represent only a small portion of the total population.

The non-stationary model with mutation rate 1E-5 at 2.5E4 replications shows the wild-type variant still at very close to a relative frequency of 1, the mutant variant at the relative frequency of about 5E-5, and the expected number of 1 drug-resistant mutant variant in that population. When the mutation rate is lowered to 1E-9, at 2E8 replications still shows a relative frequency of the wild-type to be almost 1, and the relative frequency of the mutated variant is still less than 1E-8, but one would expect there would be one mutated drug-resistant variant in that population. This is demonstrated by the Kishony Mega-Plate experiment. The vast majority of the members in a given colony are not mutated variants that can grow in the next higher drug region, these members are clones of the founder of that colony (at least at that particular site in the genome).

If the mutations in any evolutionary Markov process are accumulated sequentially as in the Kishony Mega-Plate experiment, the number of replications required to make the transition is exponentially smaller than when the mutations must be accumulated simultaneously. The mathematical explanation for this is the multiplication rule of probabilities. This is the reason that the Kishony Mega-Plate experiment can only operate with small increases in drug-concentration on the plate used. Any higher concentration of the drug or the use of two or more drugs will require a much, much larger plate to accommodate the much larger colony size necessary for such an evolutionary process.

There are two significant points to consider when a single drug is used versus two drugs (or a higher drug-concentration in the adjacent region) in the physical behavior and mathematical modeling of the Kishony Mega-Plate experiment. When an evolutionary process such as the one drug experiment is carried out, the mutations can accumulate sequentially, one at a time. What that means biologically is that some member of the population gets the beneficial mutation that gives improved fitness in a particular colony (lineage). That member is then able to form a new colony in the next higher drug-concentration region and start a new Markov process. It doesn’t matter what happens to its progenitor colony. It can continue to grow or go extinct. In the case where two drugs are used or when the drug-concentration in the adjacent band is too large (requiring more than one mutation to grow in that region), a second or higher-order Markov transition matrix is required. What this means numerically and biologically is that the variant with one of the beneficial mutations (after several hundred million replications in that colony) must continue to grow in that colony. That variant with the first beneficial mutation will have to do about 30 doublings (when the mutation rate is 1E-9) to get the second beneficial mutation necessary to grow in the next higher drug-concentration region. But the other variants will also be doubling at the same time instead of starting with just one member in that lineage it will be one of the other hundreds of millions of other members in that colony. This means the carrying capacity of that environment must be vastly larger to accommodate this much, much larger colony.

The importance of understanding DNA evolution cannot be overstated. The impact on the health care system of infectious diseases and the evolution of drug resistance is one of the greatest burdens on the medical system. According to the Healthcare Cost and Utilization Project (HCUP) [19] on the most common medical reasons in 2003 for all hospitalizations that began in the emergency department, pneumonia was the number 1, urinary infections 12, skin infections 15, and sepsis 16. These statistics have not improved with time. According to recent HCUP data (2018) [20], excluding maternal and neonatal stays, the main reasons for hospital admission are, septicemia number 1 and pneumonia number 4. That is only half the story. HCUP 2008 data [21] for the most expensive hospitalization shows the septicemia is the number 1 most expensive cause for hospitalization.

How much of this problem is due to the way antibiotics are used in the outpatient environment is unclear. Conflicting signals are give to primary care physicians on the use of antibiotics. Primary care physicians are being warned in the overuse of antibiotics because of the selection of drug resistant variants and killing non-pathogenic bacteria. [22–24] On the other hand, the data from the previous paragraph would seem to indicate that delay or under-use of antibiotics may be occurring. Primary care physicians usually don’t have access to stat laboratories to give objective evidence for a disease they are trying to treat in an ill patient. Primary care physicians must depend on the medical history and clinical examination with minimal data, usually just the patient’s vital signs and perhaps a rapid in office test such as a rapid group A streptococcal or influenza test. Physicians working in the outpatient environment don’t have the benefit of close patient observation such as what occurs with the hospitalized patient. Outpatient physicians must depend upon the patient or family members to report on condition changes and they may not be capable to recognize a worsening condition. In the case of early sepsis, even a 24 hour delay in the initiation of antibiotics can lead to life threatening infections.

The discussion does not end here. Drug-resistant infections are a bigger problem in the hospitalized patient than in the out-patient environment. It is well known that community acquired MRSA infections are still sensitive to more antibiotics than hospital acquired MRSA infections. [25] This is most likely due to the fact that hospitalized patients tend to be sicker with weaker immune systems than the generally more healthy out-patients. [26]

Based on the analysis of the evolution of drug-resistance in the Kishony Mega-Plate experiment, it is clear that the evolution of drug-resistance to a single drug selection pressure is much easier for a bacterial population to accomplish than evolution to two or more drugs simultaneously. This would seem to point to a better solution of using combination antibiotics rather than not using antibiotics to prevent the selection of resistant variants.

DNA evolution can be modeled as a Markov process, but the assumption that this Markov process is stationary leads to inaccurate predictions when doing phylogenetic DNA analysis, or using these models to predict the behavior of evolutionary experiments. The underlying problem with the Jukes-Cantor and derivative models is that in the formulation of these models, it is assumed that the evolutionary process is stationary, and these models don’t take into account population size. Implicit in the derivation of the Jukes-Cantor and derivative models is that the substitution matrix does not change. The probabilities in the transition matrix are a function of population size. DNA evolution is not a stationary Markov process, and applying this stationary model selectively to only homologous portions of the genome ignores all the genetic differences that would defeat the accuracy of the predictions this model is capable of doing. By including the population size in the transition matrix, this model will correctly simulate and predict the evolutionary behavior of the Kishony Mega-Plate experiment.

The evolution of drug-resistance of microbes to drug therapies or cancers to targeted therapies requires an accurate understanding of the evolutionary process. The non-stationary Markov chain model of DNA evolution gives an important tool for understanding evolution. And that tool gives the ability to estimate the number of selection pressures (antibiotics or targeted cancer therapies) necessary to address and suppress the evolutionary process and have a greater probability of having treatment success.

## Statements

### Statement of Ethics

No human or animal studies were used in this research. No <u>study approval statement</u> or <u>consent to participate statement</u> required.

### Author Contributions

Alan Kleinman, MD, PhD is the sole author of this paper.

### Data Availability Statement

All data and FORTRAN computer programs used to generate data are available on request in the supplemental documentation.

## References

1. Baym M, Lieberman TD, Kelsic ED, et al. Spatiotemporal microbial evolution on antibiotic landscapes. Science. 2016;353(6304):1147–1151. doi:10.1126/science.aag0822 https://www.ncbi.nlm.nih.gov/pmc/articles/PMC5534434/

2. Baym M, Gross R, How To Make A MEGA-plate, https://openwetware.org/wiki/How_To_Make_A_MEGA-plate

3. Kishony R, EXTRA MINUTES - SUPERBBUGS (Harvard Experiment explained), https://www.youtube.com/watch?v=Irnc6w_Gsas&t=1s

4. Jukes, T.H. and Cantor, C.R. (1969) Evolution of Protein Molecules. In: Munro, H.N., Ed., Mammalian Protein Metabolism, Academic Press, New York, 21–132. http://dxdoi.org/10.1016/B978-1-4832-3211-9.50009-7

5. Kimura M., “A simple method for estimating evolutionary rates of base substitutions through comparative studies of nucleotide sequences”. Journal of Molecular Evolution. 16 (2): 111–20. doi:10.1007/BF01731581.

6. Kimura M., “Estimation of evolutionary distances between homologous nucleotide sequences”. Proceedings of the National Academy of Sciences of the United States of America. 78 (1): 454–8. doi:10.1073/pnas.78.1.454.

7. Felsenstein J., “Evolutionary trees from DNA sequences: a maximum likelihood approach”. Journal of Molecular Evolution. 17 (6): 368–76. doi:10.1007/BF01734359.

8. Hasegawa M, Kishino H, Yano T., “Dating of the human-ape splitting by a molecular clock of mitochondrial DNA”. Journal of Molecular Evolution. 22 (2): 160–74. doi:10.1007/BF02101694.

9. Tamura K., “Estimation of the number of nucleotide substitutions when there are strong transition-transversion and G+C-content biases”. Molecular Biology and Evolution. 9 (4): 678–87. doi:10.1093/oxfordjournals.molbev.a040752.

10. Tamura K, Nei M (May 1993). “Estimation of the number of nucleotide substitutions in the control region of mitochondrial DNA in humans and chimpanzees”. Molecular Biology and Evolution. 10 (3): 512–26. doi:10.1093/oxfordjournals.molbev.a040023.

11. Casanellas M., Fernández-Sánchez J., Garrote-López, M., “Distance to the stochastic part of phylogenetic varieties”, Journal of Symbolic Computation, Available online 22 September 2020, © 2020 Elsevier Ltd, doi:10.1016/j.jsc.2020.09.003

12. Karcher MD, Carvalho LM, Suchard MA, Dudas G, Minin VN (2020) Estimating effective population size changes from preferentially sampled genetic sequences. PLoS Comput Biol 16(10): e1007774. https://doi.org/10.1371/journal.pcbi.1007774

13. Vavilova, V., Konopatskaia, I., Blinov, A. et al. Genetic variability of spelt factor gene in Triticum and Aegilops species. BMC Plant Biol 20, 310 (2020). https://doi.org/10.1186/s12870-020-02536-8

14. Chong, E.T.J., Neoh, J.W.F., Lau, T.Y. et al. Genetic diversity of circumsporozoite protein in Plasmodium knowlesi isolates from Malaysian Borneo and Peninsular Malaysia. Malar J 19, 377 (2020). https://doi.org/10.1186/s12936-020-03451-x

15. Maloney EM, …, Esquivel CO, Martinez OM, “Genomic variations in EBNA3C of EBV associate with posttransplant lymphoproliferative disorder”, JCI Insight. 2020;5(6):e131644. https://doi.org/10.1172/jci.insight.131644.

16. Huelsenbeck JP, Larget B, Alfaro MR, Bayesian Phylogenetic Model Selection Using Reversible Jump Markov Chain Monte Carlo, Molecular Biology and Evolution, Volume 21, Issue 6, June 2004, Pages 1123–1133, https://doi.org/10.1093/molbev/msh123

17. Wikipedia, Markov Chain, https://en.wikipedia.org/wiki/Markov_chain

18. Personal email communication, Kishony R.

19. Elixhauser A, Owens P. Reasons for Being Admitted to the Hospital through the Emergency Department, 2003: Statistical Brief #2. 2006 Feb. In: Healthcare Cost and Utilization Project (HCUP) Statistical Briefs [Internet]. Rockville (MD): Agency for Healthcare Research and Quality (US); 2006 Feb-. Available from: https://www.ncbi.nlm.nih.gov/books/NBK63506/

20. Agency for Healthcare Researcher and Quality, HCUP Fast Stats - Most Common Diagnoses for Inpatient Stays, https://www.hcup-us.ahrq.gov/faststats/NationalDiagnosesServlet

21. Friedman B, Henke RM, Wier LM. Most Expensive Hospitalizations, 2008: Statistical Brief #97. 2010 Oct. In: Healthcare Cost and Utilization Project (HCUP) Statistical Briefs [Internet]. Rockville (MD): Agency for Healthcare Research and Quality (US); 2006 Feb-. Available from: https://www.ncbi.nlm.nih.gov/books/NBK52654/

22. Ventola CL. The antibiotic resistance crisis: part 1: causes and threats. P T. 2015 Apr;40(4):277–83. PMID: 25859123; PMCID: PMC4378521.

23. Llor, C., & Bjerrum, L. (2014). Antimicrobial resistance: risk associated with antibiotic overuse and initiatives to reduce the problem. Therapeutic advances in drug safety, 5(6), 229–241. https://doi.org/10.1177/2042098614554919

24. Chang Y, Chusri S, Sangţhong R, McNeil E, Hu J, Du W, Li D, Fan X, Zhou H, Chongsuvivatwong V, Tang L. Clinical pattern of antibiotic overuse and misuse in primary healthcare hospitals in the southwest of China. PLoS One. 2019 Jun 26;14(6):e0214779. doi:10.1371/journal.pone.0214779. PMID: 31242185; PMCID: PMC6594576.

25. Samanthi, Difference Between HA-MRSA and CA-MRSA, Posted Nov. 22, 2017, https://www.differencebetween.com/difference-between-ha-mrsa-and-vs-ca-mrsa/

26. Margolis E, Rosch JW. Fitness Landscape of the Immune Compromised Favors the Emergence of Antibiotic Resistance. ACS Infect Dis. 2018 Sep 14;4(9):1275–1277. doi:10.1021/acsinfecdis.8b00158. Epub 2018 Aug 2. PMID: 30070470; PMCID: PMC6358436.

